# Highly Multiplexed Single-Cell RNA-seq for Defining Cell Population and Transcriptional Spaces

**DOI:** 10.1101/315333

**Authors:** Jase Gehring, Jong Hwee Park, Sisi Chen, Matthew Thomson, Lior Pachter

## Abstract

We describe a universal sample multiplexing method for single-cell RNA-seq in which cells are chemically labeled with identifying DNA oligonucleotides. Analysis of a 96-plex perturbation experiment revealed changes in cell population structure and transcriptional states that cannot be discerned from bulk measurements, establishing a cost effective means to survey cell populations from large experiments and clinical samples with the depth and resolution of single-cell RNA-seq.

---

Massively parallelized single-cell RNA-sequencing (scRNA-seq) is transforming our view of complex tissues and yielding new insights into functional states of heterogeneous cell populations. Currently, individual scRNA-seq experiments can routinely probe the transcriptomes of more than ten thousand cells^1,2^, and in the past year the first datasets approaching and exceeding one million cells have been reported^3,4^. However, despite numerous technical breakthroughs that have increased cell capacity of many scRNA-seq platforms, researchers are at present limited in the number of samples that can be assayed. Many biological and therapeutic problems rely on finding genes or signals responsible for a phenotype of interest, but the enormous space of possible variables calls for screening hundreds, or even thousands, of conditions. At present, analyzing genetic, signaling, and drug perturbations (and their combinations) at scale with scRNA-seq is impeded by microfluidic device operation, high reagent costs, and batch effect. While a multiplexing method based on epitope expression has been developed^5,6^, it can only be practically applied to about a dozen samples. The *in silico* demuxlet algorithm^7^ is more scalable but requires samples from distinct genetic backgrounds.

The scRNA-seq sample multiplexing method presented here allows for cells from individual samples to be rapidly chemically labeled with identifying DNA oligonucleotides, or sample tags (**Fig. 1**). This universal approach can be applied to cells from any organism without the need for specific epitopes, sequence markers, or genetic manipulation, and is compatible with any scRNA-seq protocol based on poly(A) capture. We demonstrate the utility and versatility of our technique in the context of a multifaceted experimental perturbation in which neural stem cells (NSCs) were exposed to 96 unique combinations of growth factors, with the perturbed cell populations profiled as a single pooled library (**Fig. 1a**). Despite the cell capacity of scRNA-seq platforms, single-cell transcriptome-wide analysis of such an experiment, which produces a unique cell population in each condition, has been technically and financially inaccessible in the absence of a suitable means of sample pooling. This experiment introduces a powerful new experimental and analytical paradigm, underpinned by our flexible, scalable cell tagging procedure, in which the massive cell capacity of scRNA-seq is effectively leveraged to analyze and compare large numbers of cell populations.

**Figure 1.**
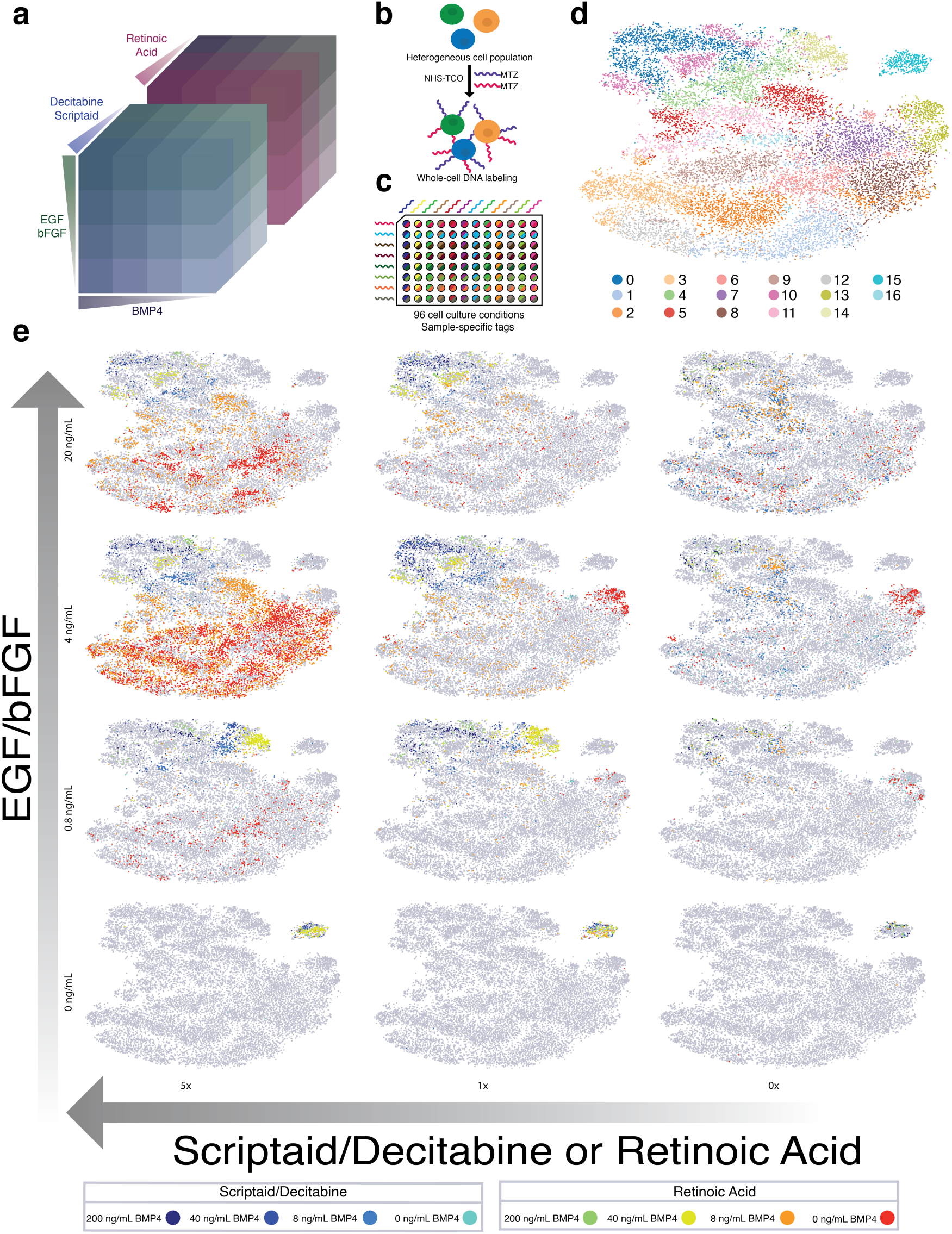
96-Plex scRNAseq experiment (**a**) Four experimental factors (EGF/bFGF, BMP-4, Decitabine/Scriptaid, and Retinoic acid) were titrated against one another to produce an array of 96 unique perturbations. (**b**) Prior to scRNA-seq, a one-pot, two-step reaction with MTZ-DNA and NHS-TCO labeled cells with sample-specific tags (**c**) Neural stem cells subjected to a 96-plex array of growth conditions were dual-labeled with a unique pair of sample tags (**d**) t-SNE of 21,232 cells from 96-plex perturbation. Cluster assignments closely match population behavior driven by experimental parameters (**e**) Visualization of cell populations produced by each experimental condition. Each t-SNE corresponds to a given EGF/bFGF concentration against a series of retinoic acid or Scriptaid/Decitabine concentrations and displays eight samples colored by BMP-4 concentration.

Neural stem cells (NSCs) are known to differentiate into many unique cell types in vivo, primarily neurons, astrocytes, and oligodendrocytes^8^. In vitro, NSCs can be forced into different differentiation trajectories by exposing the cells to a variety of synthetic chemicals, hormones, and growth factors. We investigated the response of NSCs to varying concentrations of Scriptaid/Decitabine, epidermal growth factor (EGF)/basic fibroblast growth factor (bFGF), retinoic acid, and bone morphogenic protein 4 (BMP4), producing a 4×4×6 perturbation array representing a large space of experimental conditions (**Fig. 1a**). NSCs were cultured in a single 96-well plate with each sample corresponding to a unique combination of factors (**Fig. 1c, Supplementary Fig. 5**). After chemical DNA labeling (**Fig. 1b**), the samples were pooled and subjected to a modified version of the 10x Genomics Single-Cell Expression protocol. A total of 21,232 cells were detected based on cDNA counts, and sample assignment was performed for the detected cells based on the sample tags with the highest UMI counts.

Visualization of the cell populations produced by each experimental condition revealed a complex interplay between the perturbations used in this 96-plex experimental space (**Fig. 1e**). On a global level, cell proliferation varied widely across the experiment, revealing growth rates specific not just to each condition but also to each cell state across the experiment. Highly proliferative states (clusters 1, 2, 3, 6, 7, 8, 9, 12, and 16), which account for large regions of the cell state space when plotted according to t-SNE, differentially express various genes associated with cell growth and the cell cycle, including ribosomal, cytoskeletal, and cyclin-dependent proteins (**Supplementary Table 1**). Conversely, samples deprived of EGF/bFGF exhibited apoptotic phenotypes including low cell counts and expression of stress response genes such as Cryab, Mt1, and Gpx4. We sought to define the cell states produced by the array of experimental conditions, a frequently challenging procedure in scRNA-seq analysis and a potential roadblock to perturbation experiments where the presence of classical marker genes may depend on experimental conditions. Identification of functional cell states was greatly aided by the large number of samples in our experimental perturbation. Various distinct regions of transcriptome space were repeatedly populated by cells originating from multiple samples in localized regions of perturbation space, forming natural groupings of cells that were validated and assigned by clustering using Louvain community detection (**Fig. 1d**).

Plotting the cluster occupancy of each sample revealed the structure of the cell populations produced across the experiment (**Fig. 2a**). Overall trends, such as high proliferation under low BMP4 conditions and high cluster specificity under high BMP4 conditions, are readily observed. Principal component analysis of the relative cluster abundance x sample matrix was used to identify relationships between the experimental inputs (**Fig. 2b**). The experimental perturbations associate directly with the cell populations observed in the scRNA-seq samples. The absence of EGF/bFGF has a drastic effect, yielding an isolated group of samples, while BMP4 concentration has a graded effect and a strong interaction with either Scriptaid/Decitabine or retinoic acid, each of which produces a separate branch of samples when combined with the two highest BMP4 concentrations. This analysis demonstrates that multiplexed scRNA-seq can be used to classify cell populations and interpret the conditions that produced them. In the context of a perturbation experiment, relevant features of the experimental space can be learned, e.g. the strong effect of BMP4 concentration shown here. Of perhaps greater interest would be to extend this proof-of-principle to biomedical diagnostics: by applying Bayes Rule to the relative cluster abundance x samples matrix, it should be possible to infer disease conditions from high-resolution cell population observations.

**Figure 2.**
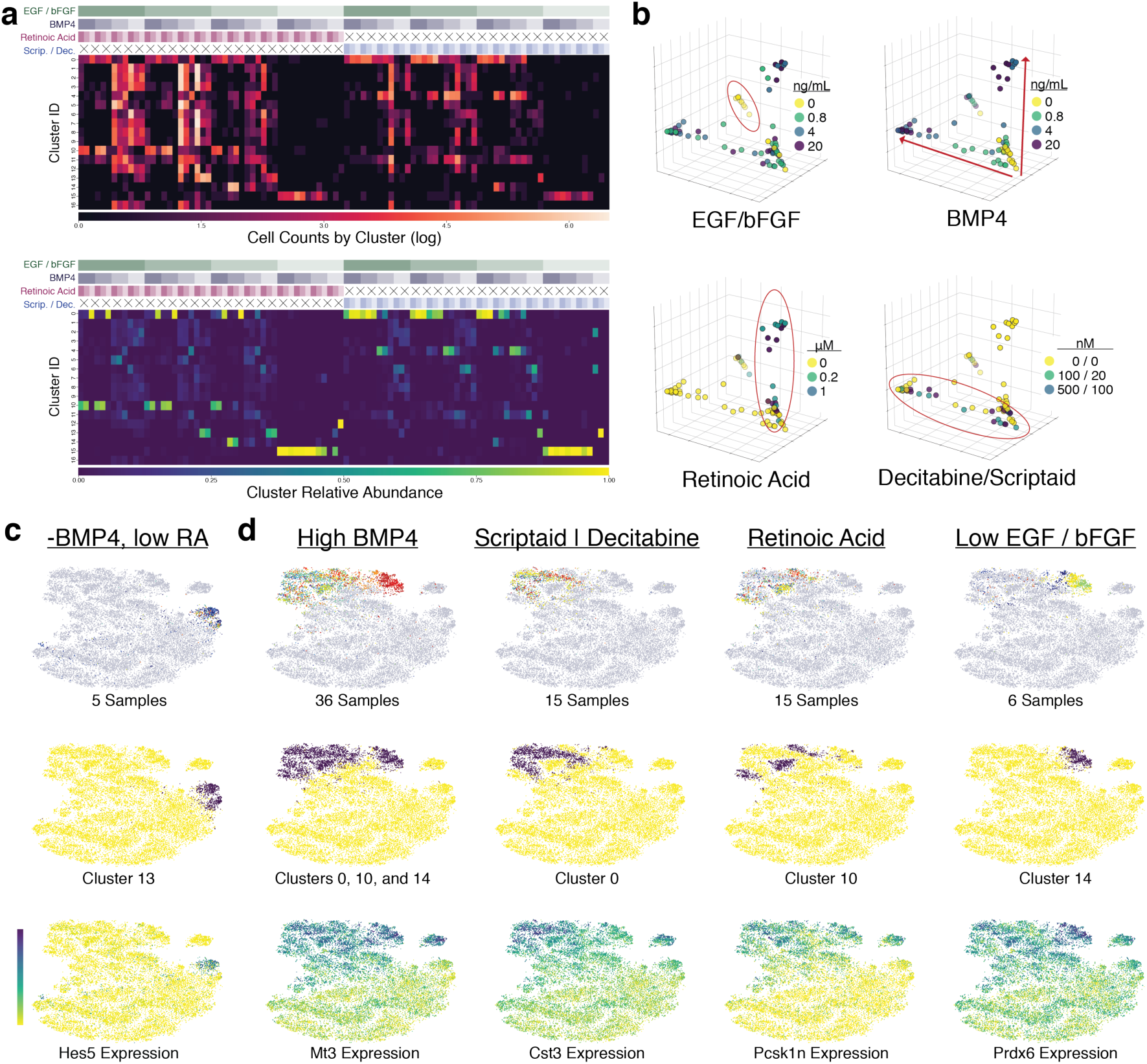
Cellular response to perturbation (**a**) Cluster occupancy versus experimental condition (above), shown as number of cells detected (top) or the relative abundance of cells assigned to each cluster in each sample (bottom). (**b**) PCA of relative cluster abundance matrix from **Fig 2a**. Each point represents a cell population from one of 96 experimental conditions, revealing patterns of influence for each experimental factor (highlighted). (**c**) Five conditions map specifically to cluster 15, characterized by Hes5 expression. (**d**) 36 samples from high BMP4 conditions map to clusters 0, 10, and 14, each specific to localized regions of the perturbation space. Expression of top differentially expressed genes for each selected group is shown.

After evaluating the high-level information that can be gleaned from a large perturbation array, we closely examined two regions of our experimental space to illustrate the depth of analysis afforded by multiplexed scRNA-seq. First, we explored an isolated portion of cell state space, cluster 13, which was populated under a strict range of conditions with intermediate EGF/bFGF concentrations, no BMP4, and moderate to no retinoic acid. Cells from just five samples accounted for practically all the cells in cluster 13 and little across the rest of cell state space, exhibiting strong condition dependence (**Fig. 2c**). Differential expression analysis showed that this cluster is strongly enriched for Hes5, a gene with important roles in cell fate determination^9^.

A more complex cellular response was observed under high BMP4 conditions, where numerous cell states were identified, many populated only within a small region of experimental space. Cells from conditions with ≥ 0.8 ng/mL EGF/bFGF and BMP4 ≥ 4 ng/mL, 36 samples in total, mapped to just three clusters (0, 10, and 14) which were further subdivided by orthogonal experimental factors (**Fig. 2d**). The cell state defined by cluster 14 was not observed in samples with high EGF/bFGF, high BMP4, or high Scriptaid/Decitabine or retinoic acid concentrations. Instead, cells from those conditions were found in clusters 0 and 10, with cells treated with Scriptaid/Decitabine appearing almost exclusively in cluster 0, while those treated with retinoic acid mapped strongly to cluster 10 with secondary populations mapping to cluster 0.

Such a dissection of cellular response to perturbations has been a long-standing goal in cell biology^11,12,13,14,15^. It has been hypothesized that cells occupy a relatively limited number of transcriptional states in response to disease or experimental perturbation, and elucidating the connections between various perturbations will help in understanding cellular behavior. One such endeavor, the Connectivity Map^14^ (CMap) project, is a large-scale effort to measure gene expression response to molecular perturbations. While impressive in scope – CMap has been used to profile more than a million perturbation experiments – major challenges have included batch effects, averaging across cell populations, and difficulty in examining conditions that yield very few cells. The multiplexing method presented here overcomes these obstacles and provides single-cell whole-transciptome resolution at very low cost.

To further validate sample multiplexing and explore its limits, we performed a multiplexing experiment in which four samples of live mouse neural stem cells (NSCs) and four samples of methanol-fixed NSCs were each labeled with unique sets of two methyltetrazine-modified sample tags. The samples were then quenched, pooled, and processed with the 10x Genomics Single-Cell Gene Expression Kit. Analysis of sample tag profiles from methanol-fixed cells recapitulated matched pairs of sample tags, indicating efficient single-cell labeling, and permitting facile sample demultiplexing (**Supplementary Fig. 2**). Cell doublet events were unambiguously detected as collisions of four pairs of tags corresponding to two separate samples. In methanol-labeled samples, we noted a strong correlation between UMI counts for pairs of tags applied to the same samples (**Supplementary Fig. 2c**), suggesting that the extent of chemical tagging may be correlated with cell size. To test this hypothesis, we devised a species-mixing experiment in which large, human HEK293T cells and small, mouse NSCs were reacted individually and in combination with a series of non-overlapping sample tag pools of increasing size (**Supplementary Fig. 3**). We found that up to 5 cell tags could be deposited on a single cell without loss of tag recovery, implying that 15,504 experiments could be multiplexed with a panel of just 20 tags. In addition, a strong correlation was observed between species of origin and sample tag counts, indicating our chemical tagging method is indeed sensitive to cell size, a relatively unexplored biological phenotype with intriguing implications for future work. Live cell labeling in aqueous solution resulted in diminished signal-to-noise (data not shown), likely a result of the high rate of NHS-ester hydrolysis in aqueous solution, along with the reduced rate of IEDDA reactions in water compared to methanol. Under methanol fixation conditions, cell tagging is a robust and flexible method for multiplex scRNA-seq with high capacity for tag multiplexing on individual cells. Compared to labeling strategies based on antibody-oligo conjugates^5,6,16^, our chemical tagging procedures are cheaper, not reliant on epitope markers, compatible with fixed cells, and, most notably, subject to chemical quenching, permitting high-throughput scRNA-seq analysis of low-input samples by pooling many cell populations before washing. While we have demonstrated multiplexing on the 10x Chromium system, our method is compatible with other similar platforms (e.g. Drop-Seq^17^, inDrops^18^, sci-RNA-seq^19^, Bio-Rad’s ddSEQ), and should be readily extendible to full-length scRNA-seq^20^ and other single-cell genomic assays.

We envision our chemical multiplexing strategy playing a central role as sequencing-based single-cell profiling continues its phenomenal increase in scale. As multiplexing of DNA libraries has vastly improved the utility and adoption of high-throughput DNA sequencing, our solution for scRNA-seq will similarly reduce costs, drive increases in cell capacity, and extend the scope of scRNA-seq beyond bulk tissue profiling. Furthermore, the increasing throughput of scRNA-seq will facilitate even higher multiplexing, and our method can be readily applied to thousands of samples. For diagnostic purposes, the cost savings associated with multiplex scRNA-seq also have the potential to accelerate the adoption of single-cell genomics in the clinic.

## Methods

### Overview of Cell Tagging Procedure

Barcoded DNA oligonucleotides (“tags”) are attached to exposed NHS-reactive amines on the cells of interest. Sample tagging is achieved in a one-pot, two-step reaction by exposing cell samples to methyltetrazine-activated DNA (MTZ-DNA) oligos and the amine-reactive cross-linker NHS-*trans*- cyclooctene (NHS-TCO) (**Fig. 1b**). NHS-functionalized oligos are formed *in situ* via inverse-electron demand Diels-Alder (IEDDA) chemistry, and nucleophilic attack by accessible cellular amines chemoprecipitates the oligos directly onto the cells. Our one-pot reaction based on the IEDDA reaction improves on a previous cell surface modification scheme^21^ that requires far higher DNA concentrations and isolation of unstable activated DNAs immediately before use. A library of methyltetrazine-modified sample tags can be prepared in advance, stored frozen for long periods, and applied to many cell samples in parallel. Sequencing library preparation is derived from recently published methods for multi-modal scRNA-seq^6,16^.

### Oligo Activation

Sample tags were prepared with either 5’- or 3’-amine modified oligonucleotides (100-250 nmol scale, Integrated DNA Technologies, **Supplementary Table 2**). HPLC purification was critical to obtain highly reactive preparations of 5’-modified oligos, while 3’-modified oligos can be purchased without HPLC purification (data not shown). In either case, oligos were resuspended to a concentration of 500 μM in 50 mM sodium borate buffer pH 8.5 (Thermo). Activation reactions were performed by combining 25 μL oligo solution with 41.8 μL DMSO (Sigma) and 8.2 μL of 10 mM NHS-methyltetrazine (Click Chemistry Tools). The reaction was allowed to proceed for 30 minutes at room temperature on a rotating platform. After 30 and 60 minutes, additional 8.2 μL aliquots of 10 mM NHS-methyltetrazine were added. After 90 minutes total reaction time, ethanol precipitation was performed by addition of 180 μL 50 mM sodium borate buffer and 30 μL 3 M NaCl. After mixing, 750 μL ice-cold ethanol was added and the mixture precipitated at −80 C overnight. The precipitate was pelleted at 20,000 x *g* for 30 minutes, washed twice with 1 mL ice-cold 70% ethanol, then resuspended in 100 μL 10 mM HEPES pH 7.2. Yield was determined by absorbance at 260 nm. Typical final concentrations ranged between 40 and 80 μM. Relative oligo activity was determined by electrophoretic mobility shift assay using Cy5-*trans*-cyclooctene (Click Chemistry Tools). Methyltetrazine-derivatized oligos were diluted 100-fold in 10 mM HEPES pH 7.2, then 4 μL of this solution was added to 1 μL of a 500 nM solution of TCO-Cy5 in DMSO. All tetrazine reactions in this work were protected from light to reduce degradation of *trans*-cyclooctene. The reaction was allowed to proceed at room temperature for 20-120 minutes and analyzed on a 12% SDS-PAGE gel. Oligo activity varied within a 2-fold range across preparations. Oligos were stored at −20 C and used without further normalization.

### Cell Culture and Fixation

Neural stem cells were cultured according to the following protocol: Cryopreserved mouse neural stem cells (NSCs) were thawed for 2 minutes at 37 °C then transferred to a 15 mL conical tube. Pre-warmed Neural Stem Cell Basal Medium (SCM003, Millipore) was slowly added to a total volume of 10 mL, and the resulting cell suspension centrifuged at room temperature for 2.5 minutes at 200 x *g*. The supernatant was removed and the cell pellet was resuspended in 2 mL pre-warmed Neural Stem Cell Basal Medium and counted on the Countess II Automated Cell Counter (Thermo). Cells were seeded on poly-L-ornithine (Millipore) and laminin (Thermo) coated 100mm culture plates at 700,000 cells per plate in 10 mL of pre-warmed Neural Stem Cell Basal Medium supplemented with EGF (Millipore) and bFGF (Millipore) at 20ng/mL each, heparin (Sigma) at 2μg/mL, and 1% Penicillin-Streptomycin (Thermo). Supplemented medium was changed the next day and every other day thereafter until confluent.

Neural stem cells for 96-sample growth factor screen were cultured according to the following protocol after previously described cell culture plate reached ~80% confluence: Stock solutions (10x) were prepared in Neural Stem Cell Basal Medium for every factor and at every concentration used: EGF+bFGF at 200ng/mL, 40ng/mL, 8ng/mL, 1.6ng/mL; BMP4 (Peprotech) at 200ng/mL, 40ng/mL, 8ng/mL, 0ng/mL; Retinoic Acid (Sigma) at 10μM, 2μM, 0 μM; Scriptaid (Selleckchem)/Decitabine (Selleckchem) at 1μM/5μM, 0.2μM/1μM, 0μM/0μM; Heparin at 20μg/mL + Penicillin-Streptomycin at 10%. 20 μL each of EGF/bFGF, BMP4, Retinoic acid or Scriptaid/Decitabine, and Heparin/Penicillin-Streptomycin were added to each well of a poly-L-ornithine and laminin coated 96-well plate for a total of 80 μL.

NSCs previously plated on 100mm culture plates until ~80% confluent were dissociated by incubation in 4 mL of ESGRO Complete Accutase (Millipore) for 2 minutes at 37 °C. After incubation, the Accutase and NSCs were transferred to a 15 mL conical tube and centrifuged at room temperature for 2.5 minutes at 200 x *g*. Supernatant was removed and the cell pellet was resuspended in 2mL Neural Stem Cell Basal Medium. Centrifugation and medium replacement were repeated one more time and cell concentration was counted on the Countess II Automated Cell Counter. The cell suspension was then diluted with additional Neural Stem Cell Basal Medium to a concentration of 18.3cells/μL. From this stock 120 μL was added to each well of the 96-well plate for a total of ~2,200 cells/well. Supplemented media for every well in the 96-well plate was replaced every other day during the 5-day incubation.

Before NSC dissociation and fixation, 80 μL of ice-cold methanol was added to each well of twelve 8-well PCR strips on an ice block. After 5 days in culture, all media in the 96-well plate were removed and the cells washed three times with 150 μL of Neural Stem Cell Basal Medium. Any remaining media were removed and replaced with 20 μL of Accutase and incubated at 37 °C for 2 minutes with gentle pipetting to help break cell clumps. Next, 20 μL of dissociated NSCs in Accutase were transferred to the 8-well strip tubes containing 80 μL of 100% methanol, and the entire volume was pipetted to mix. After fixation, the NSCs were stored at −20 °C until sample labeling.

For 4-sample NSC labeling and species-mixing experiments (below), NSCs were cultured on a 100mm poly-L-ornithine and laminin coated culture plate according to the protocol previously described until ~80% confluent. NSCs were dissociated by removing culture medium followed by incubation with 4mL Accutase for 2 minutes. NSCs in Accutase were transferred to a 15 mL conical tube and centrifuged at room temperature for 2.5 minutes at 200 x *g*. The supernatant was removed and the cell pellet was resuspended in 2mL Hank’s Balanced Salt Solution (HBSS, Thermo) with 0.04% BSA (Sigma). Centrifugation and medium replacement were repeated once and cell concentration was determined on a Countess II Automated Cell Counter. Cells were then fixed by addition of 4 volumes ice-cold methanol added slowly with constant mixing. Fixed cells were stored at −20 °C until sample labeling and scRNA-seq.

Frozen stocks of HEK293T cells (ATCC) were thawed for 2 minutes at 37 °C with gentle agitation. Thawed cells (500 μL) were added to 5 mL pre-warmed media (DMEM (Corning) + 10% FBS (Gemini Bio-Products) + 1% Penicillin-Streptomycin (Corning) and centrifuged at 1,500 x *g* for 5 minutes. The cells were resuspended in 5 mL media and transferred to a T-25 cell culture flask. Cells were grown at 37 °C with 5% CO2 following standard practices. HEK293T cells were dissociated by incubation with TrypLE Select (Thermo) for 5 minutes at 37 °C, washed twice with HBSS, and resuspended in 1 mL at a concentration of ~6 M cells/mL. Cell number and viability were measured using a Countess II Automated Cell Counter (ThermoFisher). Four mL ice-cold methanol was added slowly with constant mixing, and the resulting cell suspension incubated at −20 °C for at least 20 minutes. Cells were stored at −20 °C until sample labeling and scRNA-seq.

### Flow Cytometry and Fluorescence Microscopy

Yeast cells (Fleischmann’s Rapid Rise) were used as an abundant cellular substrate to test cell labeling reactions. Approximately 5 g of dehydrated cells were rehydrated in 4 mL PBS + 0.1%Tween-20 (Sigma) for 10 minutes at room temperature with rotation. One mL of the resulting cell suspension was diluted with 7 mL PBS-Tween and fixed by slow addition of 32 mL ice-cold methanol with constant mixing. Cells were incubated at −20 °C for at least 20 minutes before further use.

Methanol-fixed cells were rehydrated by combining 700 μL HBSS with 500 μL fixed cells in 80% methanol. This suspension was centrifuged at 3,000 x *g* for 5 minutes, then washed twice more with HBSS. Cells were resuspended in 1 mL HBSS, and 50 μL of this cell suspension was used for cell labeling. Methyltetrazine-Cy5 (Click Chemistry Tools) was added to 2 μM final concentration, NHS-TCO to 5 μM, and DAPI to 1 μg/mL. Cell labeling reactions were incubated for 30 minutes at room temperature with rotation then quenched by addition of Tris-HCl to 10 mM and methyltetrazine-DBCO (Click Chemistry Tools) to 50 μM. Samples were diluted 20-fold in HBSS and analyzed on a MACSQuant VYB flow cytometer.

Fluorescence microscopy samples were prepared as above except NHS-TCO was used at 1 μM and MTZ-Cy5 was used at 62.5 μM. Samples were imaged on a Zeiss LSM 800 laser scanning confocal microscope.

### Sample Labeling Proof of Concept

Fixed NSCs were split into four aliquots with ~400,000 cells in 100 μL 80% methanol. Live NSCs were prepared as described above, washed into HBSS, and similarly aliquoted to four samples with 400,000 cells in 100 μL. Prior to cell labeling, 8 labeling combinations were made by combing 6 μL each of two different sample tags. A 5-minute pre-incubation reaction was performed in the dark at room temperature by addition of 4 μL 1 mM NHS-TCO. After pre-incubation, cell suspensions were thoroughly mixed with the entire volume of a single sample label mix. Cell labeling proceeded for 30 minutes at room temperature on a rotating platform. Reactions were quenched by addition of Tris-HCl to 10 mM final concentration and methyltetrazine-DBCO (Click Chemistry Tools) to 50 μM final concentration. After quenching for 5 minutes, cells were pooled to create a single sample for fixed cells and a single sample for live cells. The two samples were combined with two volumes PBS-BSA and pelleted by centrifugation at 500 x *g* for 5 minutes. Cells were washed three times with PBS-BSA and vigorously resuspended in a final volume of 150 μL. Cells were analyzed and counted, then fixed and live samples were combined at equal concentration and loaded onto a single lane of the Chromium Controller (10x Genomics, Inc.) targeting 10,000 cells. Library preparation was adapted from the REAP-Seq protocol^6^. The 10x Genomics v2 Single Cell 3’ seq Reagent kit protocol (10x Genomics) was used to process samples according to the manufacturer’s procedure with modifications as follows. After initial ampilifcation of cDNA and sample tags, the two libraries were separated during SPRI size-selection. The manufacturer’s instructions were used to complete cDNA library preparation. For sample tags, rather than discarding ∼80 μL SPRI supernatant, this fraction was added to 45 μL SPRI beads and incubated at room temperature for 5 min. The SPRI beads were washed twice with 80% EtOH and sample tags eluted in 20 μL nuclease-free water. Sample tags were quantified by Qubit High-Sensitivity DNA Assay (Invitrogen) and amplified using primer R1-P5 and indexed reverse primers as appropriate (**Supplementary Table 2**). PCR was performed in a 25 μL volume including 2.5 μL sample tag library, 1.5 uL of 10 uM forward and reverse primer, 7 μL nuclease-free water, and 12.5 μL KAPA 2x HIFI PCR master mix (Kapa Biosystems). The samples were cycled as follows: 98 °C 3 min, 16 cycles of: 98 °C 20 sec, 58 °C 30 sec, and 72 °C 20 sec; and then a final extension step of 72 °C for 4 min. Final sample tag libraries were obtained using a PippinPrep automated size selection system with a 3% agarose gel set for a broad purification range from 200-250 bp (target library size is 225 bp). A Qubit assay was again used to determine library concentration for sequencing. Sample tag and cDNA libraries were analyzed on a BioAnalyzer High Sensitivity DNA kit (Agilent). Example traces are provided for reference (**Supplementary Fig. 4**). Sample tag libraries were sequenced on an Illumina MiSeq using a MiSeq V3 150 cycle kit (26×98bp reads), and cDNA libraries were sequenced on an Illumina HiSeq 4000 using a HiSeq SBS 3000/4000 SBS 300 cycle kit (2×150bp reads).

### Species Mixing and Sample Label Multiplexing

Methanol-fixed human HEK293T and mouse NSCs were prepared as described above. Samples were labeled with non-overlapping tags sets of increasing size (**Supplementary Table 3**). Suspensions of both cell types were prepared at 700,000 cells/mL in 80% methanol. Samples of 100 uL were prepared for each condition, with species mixing conditions comprising 50 μL of cell suspension from each species. For this experiment, 3’-modified oligos isolated by standard desalting were used as opposed to the 5’-modified, HPLC-purified oligos used in all other experiments presented. Tag sets were prepared by reacting 6 μL of each oligo along with 2 μL of 1 mM NHS-TCO per oligo at room temperature. After 5 minutes, the entire volume of each tag set was added to the appropriate cell suspension. Cell labeling was performed for 30 minutes at room temperature on a rotating platform. Reactions were quenched as above, pooled, and added to 2 mL PBS + 1%BSA. Samples were split across two Eppendorf tubes and centrifuged at 500 x *g* for 5 minutes. Cell pellets were resuspended in 500 μL PBS-BSA, combined, and centrifuged once more. The cell pellet was washed twice more with 1 mL PBS-BSA. Finally, the cells were resuspended in 150 μL PBS-BSA, counted, and diluted to 1×10^6^ cells/mL and loaded on a single lane of the Chromium Controller targeting 12,000 cells. Sample tag and cDNA libraries were prepared as described. Libraries were submitted as part of an Illumina NovaSeq library, targeting 500 M reads total (2×150bp reads), with sample tags submitted at 10% of the total library concentration.

### 96-Sample Growth Factor Screen

Cells for the 96-sample perturbation experiment were prepared as described above. For each sample, two sample tags (6 μL each) were combined with 4 uL 1 mM NHS-TCO according an 8×12 matrix. Columns 1- 12 of the 96-well plate correspond to tags BC21-BC32, while rows A-H correspond to tags BC33-BC40 (**Supplementary Fig. 5**). Fixed cells from each experimental condition (100 μL) were labeled with the entire volume of the corresponding sample tag mix for 30 minutes at room temperature on a rotating platform. Samples were quenched as described above, pooled, and combined with 15 mL PBS-BSA. Samples were split across two 15-mL conical tubes and spun at 500 x *g* for 5 minutes. Cell pellets were resuspended in 3 mL PBS-BSA each and centrifuged again. The pellets were washed twice with one mL PBS-BSA and resuspended in a final combined volume of 200 uL. Cells were loaded on two lanes of the 10x Chromium Controller targeting 10,000 cells per lane. Sequencing libraries were prepared as described, with sample tag libraries sequenced on two lanes of Illumina MiSeq using MiSeq v3 150 cycle kits (26×98bp reads), and cDNA libraries pooled and sequenced on Illumina HiSeq 4000 using two HiSeq 3000/4000 SBS 300 cycle kits (2×150bp reads).

### cDNA Data Processing

Standard bioinformatics tools were used to process and analyze DNA sequencing information. Raw sequencing data were processed using the 10x Genomics Cell Ranger pipeline (version 2.0). Cellranger mkfastq was used to demultiplex libraries based on sample indices and convert the barcode and read data to FASTQ files. Cellranger count was used to identify cell barcodes and align reads to mouse or human transcriptomes (mm10 and hg19) as appropriate. For the 96-sample perturbation experiment, cellranger aggr was used to combine and normalize sequencing data from the two 10x lanes split across two HiSeq lanes. Cells were selected by cellranger using the inflection point of detected cell numbers as a function of ordered read counts as a cutoff. For the sample labeling proof of concept and species mixing experiments, no further analysis of the cDNA data was performed.

### Sample Tag Data Processing and Assignment

Sequencing reads from sample tag libraries were processed using cellranger and synthetic ‘transcriptomes’ corresponding to the sequences of the tags used in a given experiment. Cellranger count outputs a post-sorted genome BAM file containing error-corrected cell barcodes and UMIs as well as read2 sequence containing sample tag information. The post-sorted genome BAM file was used to generate a digital count matrix for the sample tags corresponding to each cell barcode. A modified version of CITE-Seq Count^9^ was used to count sample tag data. Briefly, a fuzzy matching package, “fuzzywuzzy” (https://github.com/seatgeek/fuzzywuzzy), was implemented to find the sample barcode region in staggered sample tag libraries that were synthesized to improve sequencing quality. Tag reads were summed according to the combinations used in a given experiment, and sample calling was based simply on the sample with the highest number of reads. Sample assignment was performed by querying the sample tag matrix with cell barcodes identified from cDNA data, generating a vector of sample assignments that can be input into standard scRNA-seq analysis packages. For the species mixing experiment (**Supplementary Fig. 3**), in which up to five tags were used for each cell, t-SNE was performed on the sample tags x cells count matrix while *k*-means clustering was performed on a normalized count matrix in which the counts corresponding to each cell were first (1) collapsed and normalized according to the tag sets used (**Supplementary Table 3**) by adding the tag counts corresponding to each sample and dividing by the size of the tag set then (2) dividing each normalized sample count by the sum of all normalized samples for that cell.

### Data Analysis

For the 96-sample perturbation experiment, the ScanPy Python package (version 1.0.4) was used to process the filtered genes x cells matrix produced by cellranger. The data was log transformed, normalized per cell, and highly variable genes were selected as those with mean normalized counts > 0.0125 and < 3 and with dispersion > 0.5, giving 1,221 highly variable genes. The per-cell read counts were regressed out and the data scaled to unit variance. PCA was performed on this matrix, followed by t-SNE visualization based on the top 20 principal components. Clustering was performed using the neighbors and louvain tools in ScanPy with the size of the local neighborhood set to 30. For clustering based on Louvain community detection, the resolution parameter was adjusted to agree well with subpopulations produced by the perturbation experiment. We reasoned that these natural groupings represent reproducible, quantitatively distinct biological states under the conditions of our experiment and would thus hold the most information relevant to the changing experimental parameters. In practice, a resolution setting of 2 yielded clusters that agreed quite well with the sample-specific subpopulations produced by the perturbation experiment. Sample assignments were combined with cluster assignments from each cell to produce a matrix of cluster occupancy x experimental condition as well as a normalized version of the same matrix showing cluster relative abundance for each sample (**Fig. 2a**). Principal component analysis was performed on the cluster relative abundance matrix to visualize relationships between the experimental conditions used in our perturbation (**Fig. 2b**). Differential expression analysis was performed with the rank_genes_groups function in ScanPy. The top differential genes between the cluster(s) of interest and the rest of the dataset are shown (**Fig. 2c,d**).

## Acknowledgements

We thank Valentine Svensson for helpful discussion during preparation of the manuscript. Paul Rivaud assisted with the data processing workflow.

